# Chaperone-mediated Autophagy Deficiency Reprograms Cancer Metabolism Via TGFβ Signaling to drive Mesenchymal Tumor Growth

**DOI:** 10.1101/2022.07.07.499098

**Authors:** Xun Zhou, Yong Shi, Vera Shirokova, Elena Kochetkova, Tina Becirovic, Boxi Zhang, Vitaliy O. Kaminskyy, Cecilia Lindskog, Per Hydbring, Simon Ekman, Maria Genander, Erik Norberg, Helin Vakifahmetoglu-Norberg

## Abstract

The role of chaperone-mediated autophagy (CMA) in cancer initiation and progression is not well understood due to the lack of a loss-of-function cancer models of LAMP2A, the key regulator of this process. Here, by generating an isoform-specific knockout of LAMP2A, we show that CMA deficiency promotes proliferation and tumor growth in human cancers of mesenchymal origin. Accordingly, we observed that LAMP2A diminishes in metastatic lesions compared to matched primary human tumors from the same patients. Loss of CMA enhanced TGFβ signaling in tumors, rewired the tumor metabolome to promote anabolic pathways and mitochondrial metabolism, meeting the metabolic requirements of rapid growth. Mechanistically, we show that TGFβR2 enhances the enzymatic activity of glucose-6-phosphate dehydrogenase (G6PD), the rate-limiting enzyme of the pentose phosphate pathway (PPP), to promote the generation of nucleotides. Consequently, pharmacological inhibition of TGFβ-signaling in LAMP2A-KO cells suppresses G6PD activity, mitochondrial metabolism, and proliferation to WT levels. Conversely, pharmacological inhibition of mitochondrial metabolism suppressed LAMP2A-KO driven proliferation. Overall, our study provides a molecular mechanism on the CMA’s tumor-suppressive function by connecting two important oncogenic pathways, the TGFβ signaling and PPP metabolism, to the loss-of-function LAMP2A in mesenchymal cancer types.

## Introduction

During recent years, chaperone-mediated autophagy (CMA), representing one of the major autophagy pathways, is being increasingly recognized to play crucial role in various physiological and pathological conditions ^1-5^. Regarding malignancies, numerous studies have described a relationship between CMA and cancer following its first reported involvement in carcinogenicity more than 11 years ago ^6^. Ever since, both a pro-oncogenic capability of CMA in supporting cancer growth ^7, 8^ and an anti-oncogenic role, due to selective targeting of multiple notorious oncoproteins, has been described ^9-11^. However, our understanding of CMA’s role in cancer mostly relies on experiments performed on knockdown studies ^6, 9, 10, 12^ or in mice ^7^. Thus, due to the absence of CMA loss-of-function human cancer models, its role in tumor growth and progression remains consequently elusive.

A critical molecular component of CMA is the LAMP2A protein, which facilitates substrate uptake by the lysosome. To date, LAMP2A is the major determinant and rate limiting regulator of the CMA process, and its levels are considered to directly correlate with CMA activity. Yet, human LAMP2A knockout (KO) cancer models are currently lacking. Here, to evaluate CMA in cancer, we generated the first human cell isoform-specific knockout (KO) of LAMP2A, and our findings challenge the general current view on CMA in cancer. We find that the KO of LAMP2A drives mesenchymal tumor growth, suggesting that the metastatic potential of cancer cells is an important determinant in CMA’s tumor suppressive role. In support of this, we show that LAMP2A is more abundant in normal non-neoplastic tissues compared to human tumors, and its expression diminishes in metastatic tumors compared to the matched primary lesion, thus with malignant progression in the same patient. The tumor suppressive function of CMA is elicited by a reciprocal interaction with TGF-β signaling and a rewiring of the tumor metabolome, which results in growth advantage of CMA-deficient tumors.

## Results

### LAMP2A deletion promotes cell proliferation and growth of mesenchymal tumors

CMA has been reported to differ between primary cancer and non-oncogenic control cells ^6, 13-15^. However, very little is known on the role of CMA on tumor biological aspects of tumor origin, stage, and metastatic potential of cancer cells. To put this in focus, we created the first isoform-specific KO of *LAMP2A* in cancer cells of epithelial (carcinoma A549) and mesenchymal origin (sarcoma HT1080) (Extended data Fig. 1A-1D), by CRISPR-Cas9 gene-editing without disrupting other isoforms of *LAMP2* or *LAMP1* gene ^16^. *In vitro* characterization of these cells revealed an approximate doubling in growth rates and clonogenicity of the KO relative to the wildtype (WT) counterparts of HT1080, but not of A549 cancer cells (Fig. 1A-B). To assess this *in vivo* and to explore the impact of CMA loss-of-function on tumor growth, WT or LAMP2A-KO cells of the same cancer type were xenografted into BALB/c nude mice subcutaneously on both flanks of each animal (Fig. 1C). Consistent with the results from *in vitro* culture, a striking difference in tumor growth was observed, as deletion of LAMP2A (Fig. 1D and Extended data Fig. 1E-1F) resulted in a significant increase in volume and weight of HT1080 tumors, compared to the WT tumors in the same mice (Fig. 1E-1G). We observed no difference in growth of WT and LAMP2A-KO tumors of the A549 cancer (Fig. 1E-1G). In fact, LAMP2A-KO HT1080 tumors showed a higher proliferative capacity than the WT as evident from a higher grade of 5-Ethynyl-2′-deoxyuridine (*EdU*) positivity (Fig. 1H). Further, compared with the control, staining intensities with ID1 (a mesenchymal marker) antibody and its protein expression was found significantly more frequent in LAMP2A-KO HT1080 tumor cells (Extended data Fig. 1G-1H), suggesting that, unlike the epithelial A549 cancer cell model, LAMP2A displays an inhibitory role in tumorigenicity of HT1080 cancer cells of mesenchymal origin. These results indicate that tumor cell-of-origin or the metastatic potential of cancer cells may be important determinants for LAMP2A’s tumor suppressive role in cancer.

**Figure 1.**
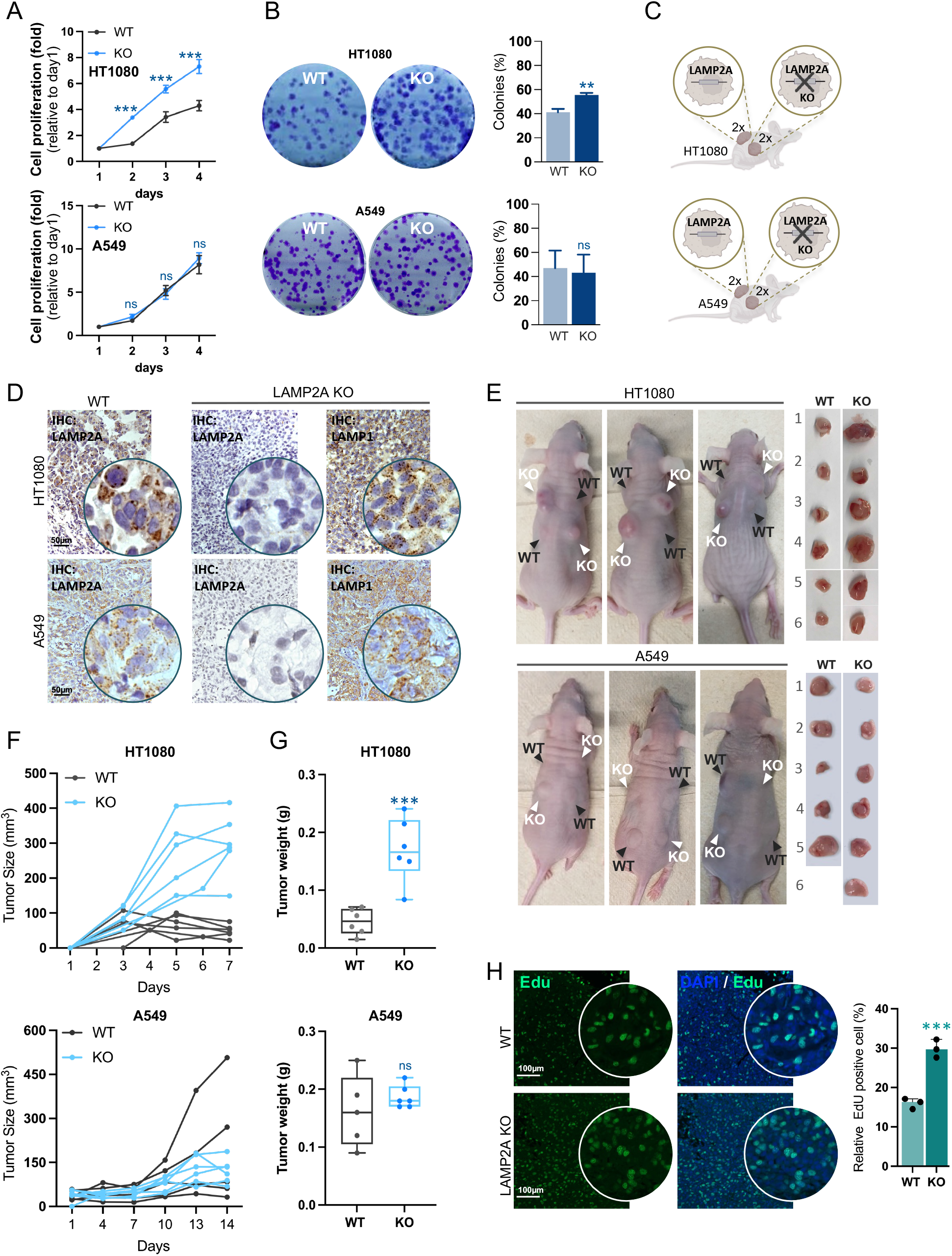
Knockout of LAMP2A drives HT1080 cell proliferation and tumor growth. The effect of genetic depletion of LAMP2A isoform on. **(A)** proliferation **(B)** clonogenic growth **(C)** experimental overview of LAMP2KO implantation of 4 tumors (2 KO vs. 2 WT) per mice. **(D)** Immunohistochemical staining for LAMP2A and LAMP1 in WT and KO tumors from HT1080 and A549 cell lines (n=3 mice). The scale bar (in black) is 50 µm. Insets at higher magnification as indicated. (E) Representative images of the xenograft mice and tumors. (F and G) Tumor growth and weight (n=3 mice). (H) Representative image of EdU positive (EdU+, Green) and DAPI-(blue) labeled HT1080 WT and KO tumor slides (n=3). Scale bar is 100 μm (H, left). Quantification of the EdU+ cell ratio per 20× field (bar graph). Error bars, ±SD. ***p<0.001; ns= non-significant; two-tailed student’s *t*-test.

### LAMP2A expression diminishes with the metastatic potential of human tumors

To determine whether CMA functionality varies along tumor staging and progression, we undertook two approaches using patient material. First, focusing on lung cancer, representing a cancer type with high metastatic potential, we probed LAMP2A levels in non-neoplastic (normal) lung tissues by RNA sequencing comparing three healthy lung tissues to eight primary lung tumors. We found that LAMP2A transcripts expressed higher in healthy tissues compared to all tested primary tumors (Fig. 2A). We then transcriptionally profiled LAMP2A in lung cancer patients with matched pairs of primary non-small cell lung cancer (NSCLC) tumors and their cognate brain metastases (Fig. 2B-2C). LAMP2A level was found to be more abundant in the primary tumors when compared to the matched metastatic tumors from the same patients (Fig. 2B). Notably, this observation was made on patient groups with primary tumors displaying epithelial traits, while their metastatic tumors showed an enrichment of mesenchymal markers (Fig. 2D), signifying that CMA is less functional in advanced cancers or actively suppressed during progression of mesenchymal tumors.

**Figure 2.**
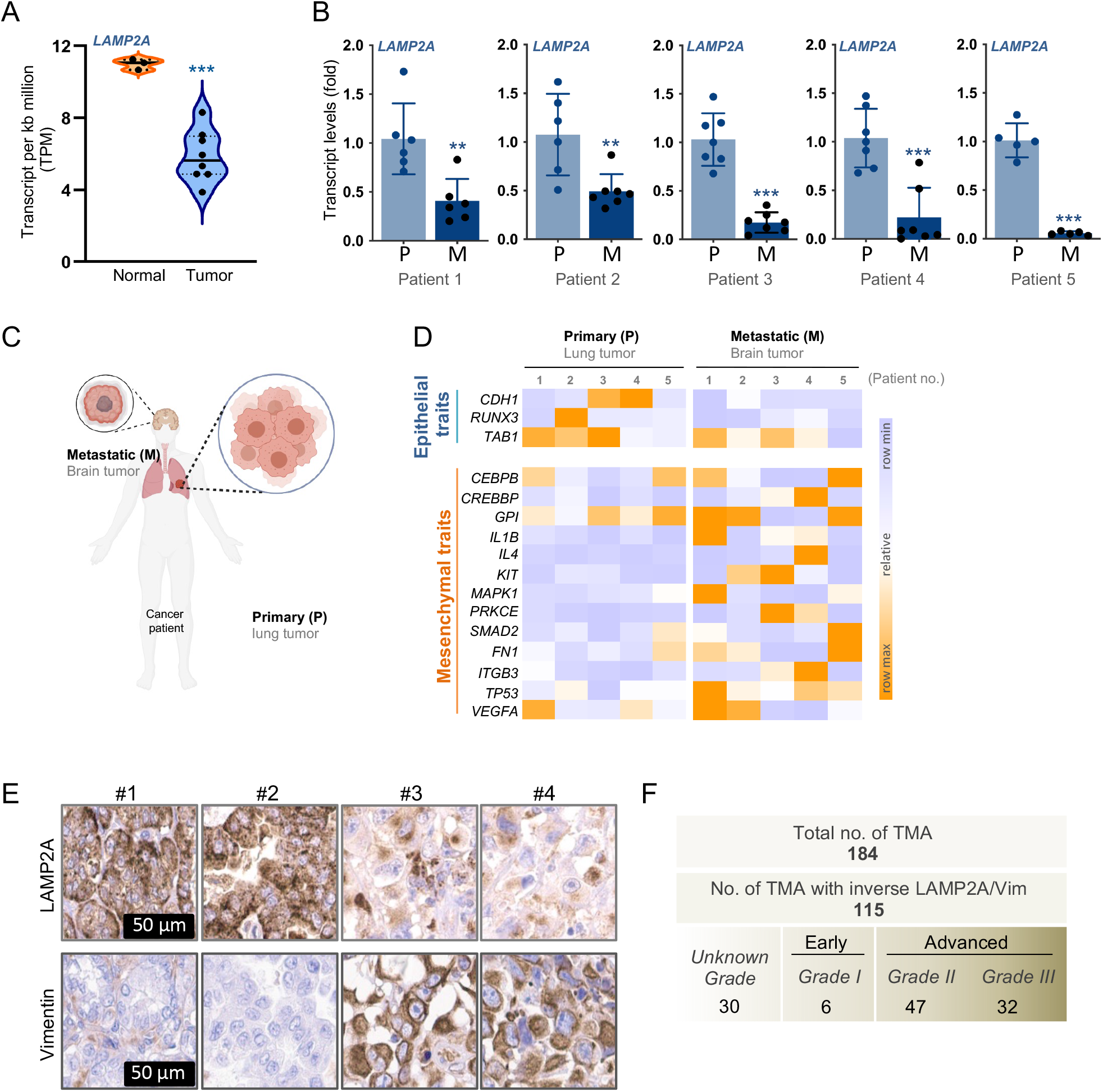
The expression of LAMP2A is lower in human metastatic lesions compared to matched primary tumors. **(A)** The transcript abundance of *LAMP2A* in three non-neoplastic human lung tissues compared to 8 primary human lung tumors. **(B)** *LAMP2A* expression in human metastatic and primary cancer tissues (n=5, for each patient). **(C)** Schematic picture of human NSCLC primary tumors and brain metastasis in cancer patient. **(D)** Comparative RNA expression analysis of human NSCLC primary tumors to match brain metastasis for EMT markers. **(E)** Tissue-Microarray analysis (TMA) of the LAMP2A and Vimentin protein expression level in 184 human multiorgan metastasis array. Two representative cores are shown in the figure to illustrate the inverse LAMP2A/Vimentin staining patterns. Scale bar 50 µm. **(F)** table revealing the number of patients with inverse expression of LAMP2A/Vimentin and its association with tumor grade. Error bars, ±SD. **p<0.01, ***p<0.001; ns= non-significant; two-tailed student’s *t*-test.

Next, we performed a large-scale LAMP2A immunohistochemical analysis on tissue arrays from metastatic cancer of different stages and lesions, encompassing 184 patients (from ovary, liver, colon, kidney, prostate, bladder, pancreas, larynx, stomach, small intestine, rectum, esophagus, uterus, cervix, breast, skin, skin malignant melanoma and thyroid). However, prior to this, we first validated anti-LAMP2A antibodies (ab18528, ab125068 and ab240018) for their sensitivity and specificity by multiple experimental approaches using WT and LAMP2A-KO human cancer cells and tumors. Our assessment revealed that all antibodies were suitable for Western Blotting (Extended data Fig. 2A). However, the ab18528 resulted in strong non-specific signal by immunofluorescence (IF) in the KO cells and immunohistochemical (IHC) staining in the KO tumors, compared to the ab125068 or ab240018 antibodies (Extended data Fig. 2B-2C). These data show that the most widely used LAMP2A antibody (ab18528) cross-reacts with an unknown target and is non-specific for the detection of this critical component of CMA in human cancer cells or tumor tissues by immunostaining applications. Taken into consideration that a caution was recently raised for LAMP2A antibody to recognize a non-specific protein in the liver of LAMP2A-KO mice ^1^, we believe that these findings raise concerns and warrant a thorough re-evaluation of LAMP2A expression, laying the groundwork for research in the CMA field.

Accordingly, we employed the anti-LAMP2A antibody ab240018, which was undetectable in the KO, for staining of histological sections at the Human Protein Atlas. Co-assessment of the expression of Vimentin with LAMP2A in 184 samples showed an inverse staining pattern in 115 (63%) of the patients (Fig. 2E). We identified that the inverse LAMP2A/Vimentin correlation was more frequently observed (69%) in metastatic lesions of advanced tumors (Fig. 2F), showing that tumors of higher grade with less LAMP2A, indicative of low CMA capability, displayed higher Vimentin levels. Since Vimentin is a classical mesenchymal marker and an oncoprotein ^17^, which confers the ability to invade and metastasize, these data illustrate a strong association of CMA deficiency with increased mesenchymal phenotype of the tumors.

Combined, these results indicate that LAMP2A expression is diminished with tumor progression and is highly correlated with the Epithelial–mesenchymal state of the tumors.

### CMA deficiency rewires the tumor metabolome

To address the reason behind the observed growth advantage of CMA deficient tumors, we reasoned that LAMP2A-KO cancer cells may be able to rewire their metabolism to meet the demands of rapid proliferation. To test this hypothesis and whether corresponding tumorigenicity of LAMP2A-KO tumors were associated with distinct metabolic patterns, we focused into multiple metabolic pathways that could contribute to the generation of biomass required for fast dividing cells (Fig. 3A). We compared the metabolomes of WT and LAMP2A-KO tumors by targeted quantitative mass spectrometry-based metabolomics approach monitoring >110 key metabolites of the major metabolic pathways. The sample groups were compared by principal component analysis (PCA), which verified a clear separation of the metabolomic profiles of the triplicates of each tumor group (Fig. 3B). A heatmap representation confirmed that the LAMP2A-KO tumors have acquired a significantly distinct metabolic profile compared to WT tumors (Fig. 3C).

**Figure 3.**
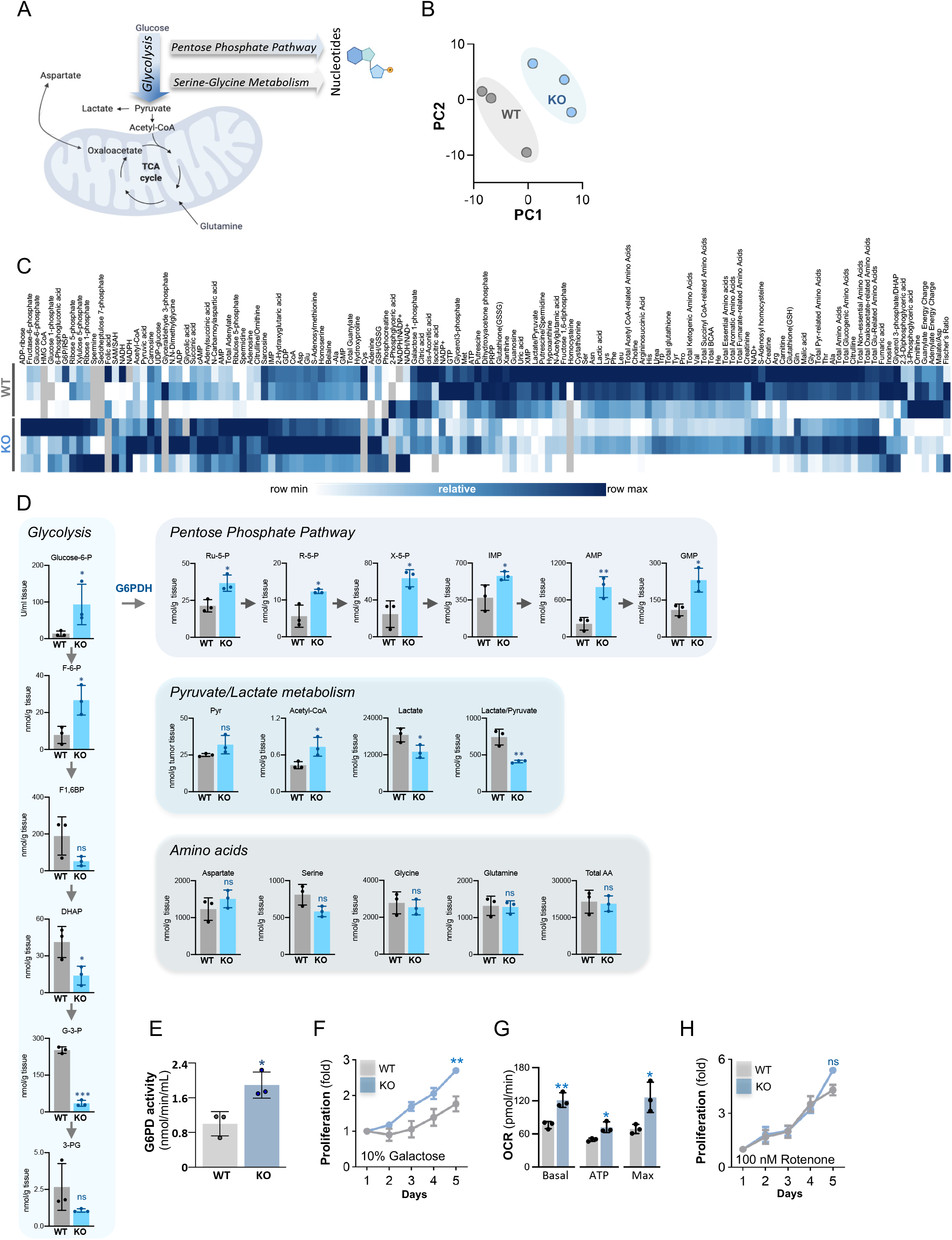
CMA Deficiency Reshapes the Tumor Metabolome to Meet the Metabolic Requirements to Support Proliferation. **(A)** Schematic picture of metabolic pathways important for nucleotide production. **(B)** Principal component analysis of the biological triplicates of the two sample groups. **(C)** Heatmap representation of the detected metabolites per biological replicate and sample group. **(D)** Individual metabolites of central metabolic pathways including glycolysis, pentose phosphate pathway, Pyruvate/Lactate metabolism and amino acids. (**E**) Enzymatic activity assay of G6PDH as nmol/min/ml. Assessment of the proliferation rate of the HT1080 WT and KO cells in media containing. **(F)** 10% galactose containing media **(G)** Oxygen consumption rate (OCR) in HT1080 WT and cells. **(H)** Proliferation of the HT1080 WT and KO cells after the treatment with 100 nM Rotenone (inhibitor of complex I of the respiratory chain). Error bars, ±SD. *p<0.05, **p<0.01, ***p<0.001 ns= non-significant; two-tailed student’s *t*-test.

Overall, we identified the metabolite levels of the pentose phosphate pathway (PPP) to be significantly more abundant in the LAMP2A-KO tumors (Fig. 3D, top panel). As nucleotide metabolism is enhanced to ensure a steady supply of building constituents required for RNA and DNA biosynthesis ^18^, their higher production represents essential component of cell division that correlates well with the proliferation rate ^19^. Correspondingly, we found the biosynthetic output of the PPP, *i*.*e*., nucleotides (IMP, AMP and GMP) to be enriched in the LAMP2A-KO tumors (Fig. 3D). The higher levels of PPP metabolites in the CMA deficient tumors raised the possibility that the pathway may be more active in LAMP2A-KO tumor as compared to WT. In line with this, measurement of the rate-limiting enzyme activity of glucose-6-phosphate dehydrogenase (G6PD), showed an approximate 2-fold increase in the LAMP2A-KO cells (Fig. 3E), indicating that higher G6PD activity accounts for the boosted PPP. These data are consistent with a recent study observing that loss of G6PD activity occurs in cancers with activated NRF2, a transcription factor reported to regulate LAMP2A levels ^20^.

Further, while the levels of the glycolytic intermediates varied, they did not differ in a consistent way between WT and LAMP2A-KO tumors (Fig. 3D, left panel). However, the end-product of glycolysis (lactate) was significantly lower (Fig. 3D) along with a deceased lactate to pyruvate ratio (Fig. 3D, middle panel), suggesting that mitochondrial metabolism may be elevated while glycolysis suppressed in the LAMP2A-KO tumors. To functionally assess the functional role of glycolysis and mitochondrial metabolism to cell growth, we performed supplementation assays (to suppress glycolysis). Galactose supplementation over 5 days reduced the growth of both WT and KO by >60% (Fig. 3F), compared to growth in normal media (Fig 1A), arguing that the glycolysis may not be the metabolic feature that specifically provides the KO tumors with a growth advantage. Instead, as Acetyl-CoA levels were elevated in the KO indicating that mitochondrial metabolism may be relevant in this context (Fig. 3E, middle panel), we measured the mitochondrial oxygen consumption rates of the cells using an Extracellular Flux Analyzer. We found that mitochondrial activity was higher in the KO compared to the WT (Fig. 3G), and importantly observed that suppression of mitochondrial metabolism by a low non-toxic concentration of rotenone (a complex I inhibitor of the electron transport chain) eliminated the growth advantage of KO cells over WT (Fig. 3H).

Moreover, amino acids are involved in key pathways that feed cancer cells to provide building blocks required for cancer cell growth. Aspartate, glutamine, serine, and glycine are among the previously reported key amino acids linked to nucleotide biosynthesis and to be required for cancer cell proliferation and tumor growth ^21-25^. However, we found no significant changes neither in the amino acid levels across the tumor sample groups nor when measuring the total amino acid levels (Figure 3D, bottom panel).

Combined, these data suggest that the major metabolic features distinguishing the LAMP2A-KO from WT tumors are the PPP and mitochondrial metabolism, both required to support a high proliferation rate. Further, consistent with the findings that metastasizing melanoma cells depend on G6PD ^26^, our findings signify that LAMP2A deficient cancer cells undergo a metabolic switch to become more reliant on G6PD and PPP function.

### Reciprocal Interaction of CMA and TGF-β signaling

To investigate the potential mechanistic details by which LAMP2A KO enhances the metabolism and tumorigenicity of cancer cells, we explored a panel of the same cancer type spanning from epithelial to mesenchymal phenotype. We found LAMP2A to be expressed less in cancer cells with enhanced mesenchymal markers, mirroring our observations in human clinical samples (Fig. 4A and Extended Fig. 3A). We specifically noted that TGF-β receptor 2 (TGFβR2) was inversely expressed at high levels in the mesenchymal cell lines to LAMP2A (Fig. 4A).

**Figure 4.**
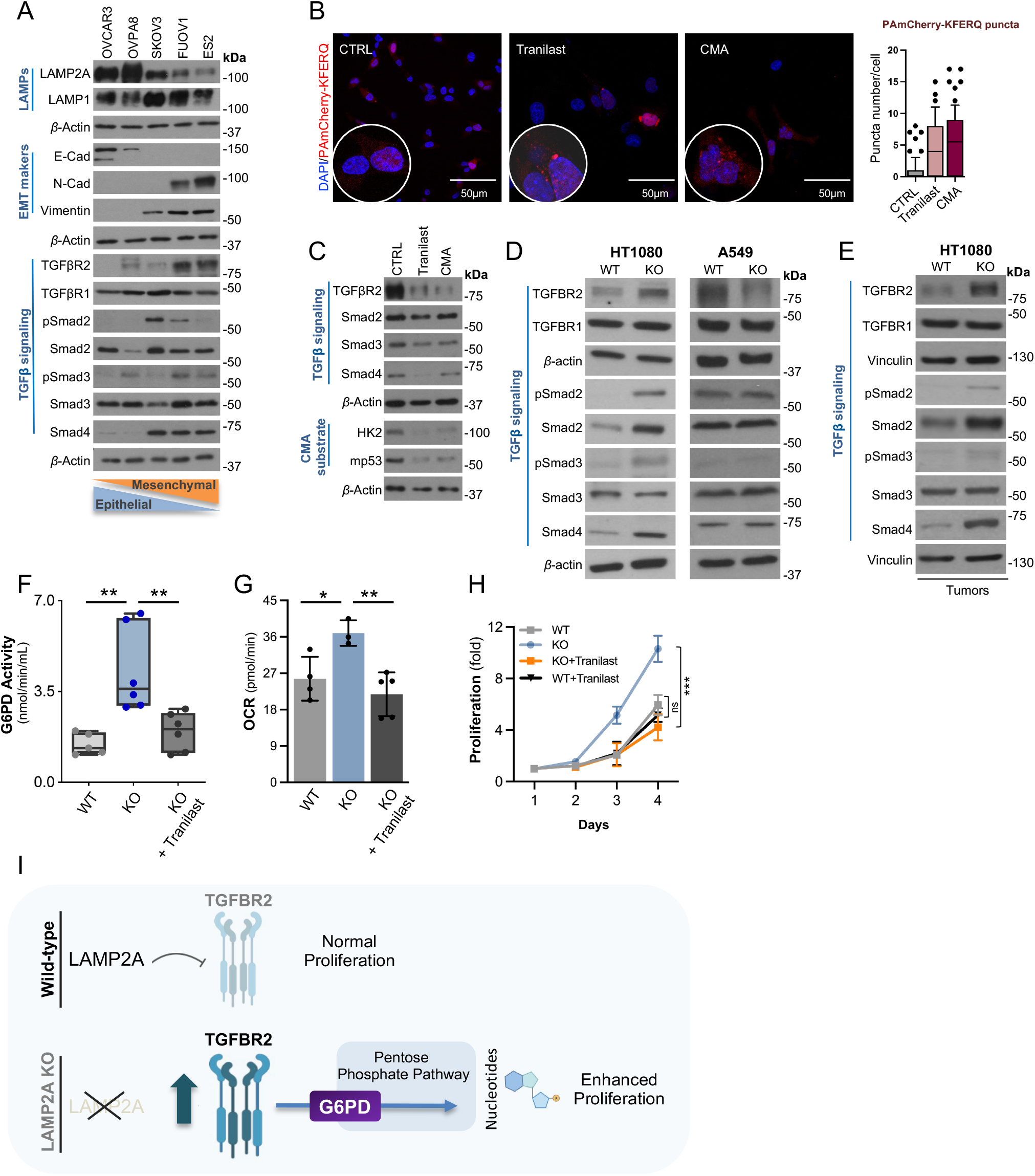
Reciprocal interaction of CMA and TGFβ signaling. **(A)** Western blot analysis of an EMT cell line panel. The effect of inhibition of TGFβ signaling by Tranilast on **(B)** CMA activity as measured by immunofluorescence (PAmcherry-KFERQ puncta) **(C)** and TGFβ signaling and degradation of CMA substrates **(D)** Western blot analysis of TGFβ signaling in HT1080 and A549 cell lines and **(E)** tumors. **(D)** CMA activity as measured by immunofluorescence (PAmcherry-KFERQ puncta) **(F)** G6PDH activity **(G)** mitochondrial respiration monitored by Seahorse Technology. **(H)** Proliferation of the HT1080 WT and KO cells after the treatment with 50 µM Tranilast. Error bars, ±SD. *p<0.05, **p<0.01, ***p<0.001; ns= non-significant; two-tailed student’s *t*-test. **(I)** Proposed working model highlighting key findings of the comparative analysis between WT and KO tumors.

As our combined findings suggest that LAMP2A diminishes upon tumor progression with increased metastatic potential, and TGF-β-induced epithelial-mesenchymal transition (EMT) is a major feature of metastasis ^27, 28^, it raised the possibility that TGF-β signaling could act to suppress LAMP2A levels. To test this hypothesis, we treated epithelial cancer cells with TGF-β1 ligand and found a lowered LAMP2A expression concomitant with decreased CDH1/E-cad and increased Vimentin levels, indicative of EMT induction (Extended Fig 3B). In contrast, we found that the knockdown of TGFβR2 or pharmacological inhibition of the TGF-β signaling with Tranilast (Rizaben) resulted in marked elevation of LAMP2A levels (Extended Fig. 3C-3D) and CMA activation (no. puncta/cell) in mesenchymal cancer cells, as measured by the CMA reporter PAmcherry-KFERQ (Fig. 4B). Tranilast treatment also caused significant reduction in TGFβR2 protein levels along with the degradation of classical CMA substrates (mutant p53 and HK2) in a comparable fashion as when CMA was activated as previously (Fig. 4C). Since both the tranilast and CMA activation-caused reduction of TGFβR2 was blunted upon chloroquine treatment (Extended Fig. 3E), along with the data showing a lower enrichment level of TGFβR2 in isolated lysosomes upon LAMP2A knockdown, determined by quantitative mass spectrometry (Extended Fig. 3F), our data suggest that CMA accounts for TGFβR2 degradation in cancer cells. In fact, the knockdown of LAMP2A in the cancer cells resulted in elevated TGFβR2 expression (Extended Fig. 3G), which was further confirmed in LAMP2A-KO HT1080 cells (Fig. 4D) and tumors (Fig. 4E), while re-expression of LAMP2A in the HT1080 LAMP2A-KO cells restored TGFβR2 levels to WT (Extended Fig. 3H). Combined, beyond revealing that mesenchymal cancer cells display less basal CMA activity, compared to epithelial cancer cells, our data show that LAMP2A levels are actively suppressed along EMT and suggest that CMA activation can be an important approach to exhaust the EMT potential of cancer cells through modulating TGF-β signaling.

Importantly, we found that inhibition of TGF-β signaling in the LAMP2A-KO cells by Tranilast could markedly restore the KO-mediated metabolic features, including the high G6PD activity (Fig. 4F) and increased mitochondrial respiration (Fig. 4G), and proliferation (Fig. 4H) down to WT levels. Thus, our study provides a molecular mechanism on the CMA’s tumor-suppressive nature by connecting two important oncogenic pathways, the TGF-β signaling and PPP metabolism, to the LAMP2A loss-of-function in mesenchymal cancer types (Fig. 4H).

## Experimental Procedures

### Human samples

The material was obtained from 5 patients with NSCLC, that underwent surgery of the primary tumor and with surgically removed brain metastases from a study that has received ethical approval by the institutional review board at Karolinska University Hospital (Registration number: 2016/944-31/1). Total RNA was isolated from formalin-fixed, paraffin-embedded (FFPE) 4 × 4 μm tissue sections of primary tumors and matched brain metastases using RNeasy FFPE kit (Qiagen, Hilden, Germany) per manufacturers’ instructions. RNA quantity and quality were assessed using RNA Screen Tapes on a 2200 TapeStation system (Agilent, Santa Clara, CA, USA). All tissue samples displayed similar RNA integrity number curves. For each sample, 200ng of RNA was submitted to Novogene Corporation Inc. (Cambridge, UK) for cDNA library construction and 150-bp paired-end sequencing using the Novaseq PE150 platform. Immunohistochemistry and slide scanning were performed as described before ^29, 30^. Multiorgan metastatic tissue arrays were purchased from Biocat. The sections were deparaffinized in xylene, hydrated in graded alcohols and blocked for endogenous peroxidase in 0.3% hydrogen peroxide diluted in 95% ethanol. A Decloaking chamber (Biocare Medical) was used for antigen retrieval. Immunohistochemical staining of vimentin, and LAMP2A was performed using an Autostainer 480 instrument (Thermo Fisher Scientific), incubating the slides with anti-vimentin (HPA001762, Atlas Antibodies Ab, diluted 1:1000), and anti-LAMP2A (Abcam, ab240018), and developed for 10 min using diaminobenzidine Quanto (Thermo Fisher Scientific) as chromogen. All incubations were followed by rinsing in wash buffer (Thermo Fisher Scientific) 2 × 5 min. Slides were counterstained in Mayers haematoxylin (Histolab) and cover slipped using Pertex (Histolab) as mounting medium. The stained slides were digitalized using the automated scanning system Leica Biosystems Deerpark, IL, USA), using a ×20 objective.

### Animal experiments

The lung cancer cell line A549 and the fibrosarcoma cell line HT1080 were obtained from ATCC. LAMP2A KO cells were generated by CRISPR-Cas9 gene-editing as described previously ^16^. All cell lines were maintained in RPMI-1640 Medium (Sigma-Aldrich, R8758) supplemented with 10% FBS (Gibco, 10500-064) and 1% penicillin-streptomycin-glutamine. All cell lines were cultured in the incubator at 37°C with 5% carbon dioxide in the air.

All animal experiments were approved by the Animal Ethical Review Board, Sweden (Ethical permit No. N116/16). All mice were maintained following the guidelines of Karolinska Institutet. For xenografting, 1 × 10^6^ cells (A549 and HT080 cells) were collected during the exponential growth phase, and xenografted subcutaneously into the flank of 7 or 8-week-old BALB/c nude female (Janvier). Total volume of one injection was 50 µl in 50% Matrigel (Corning, 356231), 4 (2×2 symmetrical) injections per mouse. Mice were sacrificed when at least one tumor had reached the maximum allowed size (10 mm^3^) as determined by calipering using the formula (3,14 x tumor length x tumor width^2^)/6 or if ulceration was observed.

Tumors were separated from the skin and accurately weighted. Then, fresh tumors were cut and the required tissues were collected for RNA isolation, immunohistochemical and immunofluorescence stainings. The remaining tissues were snap frozen in liquid nitrogen and stored at -80°C until further use.

For EdU labeling, the mice were intraperitoneal injected 10 µg/g EdU 2hrs before sacrifice. Tumor sections were prepared according to the IHC protocol l. EdU was detected by the Click-It EdU Alexa Fluor 488 Kit. All samples were imaged with a Zeiss Axio Imager M2 microscope with an Axio Cam HR camera (Zeiss). ImageJ was used for image analysis.

### Histology and microscopy

Cells were fixed for 15 min in 4 % formaldehyde and permeabilized with digitonin (0.025%, 10 min at RT). Afterwards, the cells were stained overnight (4°C) using primary antibodies (all diluted to 1:300 in 2% BSA in PBS). The next day, the samples were washed followed by staining with the secondary Alexa Fluor 594 goat anti-rabbit IgG (Invitrogen, R37117) or Alexa Fluor 488 anti-mouse IgG (Invitrogen, R37118) for 1 h at RT. Nuclei were counterstained with DAPI (1mg/ml) for 10 min at RT and slides were mounted using Vectashield mounting medium. The pictures were taken using ZEISS LSM 710 inverted confocal microscope and the 63x/1.4 oil immersion objective.

Tumors from BALB/c nude mice were fixed in 4% paraformaldehyde over night at 4°C and embedded in paraffin according to the routine procedure, 5 μm thick sections were used for staining.

Immunohistochemistry and immunofluorescence stainings were performed according to standard procedures. In brief, sections were deparaffinized followed by antigen retrieval for 10 min in sodium citrate buffer pH6.5 (Merck, C9999-100ML) in a steamer. Sections were permeabilized and blocked for 60 min in 1% BSA, 5% normal donkey serum (Jackson Immunoresearch, 017-000-121) followed by incubation with primary antibodies (all diluted to 1:500 in 1% BSA in PBS-Triton 0,1%) at 4°C overnight. The following primary antibodies were used for staining: three anti-LAMP2A antibodies (Abcam, ab18528, ab125068, ab240018). Secondary ab with the correspondent species specificity were used for the signal detection: donkey anti-rabbit Alexa Fluor Cy3 and 488 conjugated secondary ab (Invitrogen, A32790, A21207) for IF, and HRP-conjugated anti-rabbit ab (Vector Laboratories, MP-7401) followed by DAB chromogen detection (Dako, 3468) for IHC. All samples were imaged with a Zeiss Axio Imager M2 microscope with an Axio Cam HR camera (Zeiss). ImageJ was used for image analysis.

### Cell lines

The ovarian cancer cell lines OVCAR3, OVPA8, SKOV3, ES2; the fibrosarcoma cell line HT1080 and the lung cancer cell lines A549 were maintained in RPMI medium (Sigma-Aldrich, R8758) with 10% (v:v) heat-inactivated fetal bovine serum (FBS; Gibco, 10500064), 1% [v/v] penicillin/streptomycin (Sigma-Aldrich, P0781) and 1% (v:v) glutamine (Sigma-Aldrich, G7513). The ovarian cancer cell line FU-OV1 was cultured in DMEM+F12 (1:1) medium (Gibco, 11320033), supplemented with 10% (v:v) FBS and 1% [v/v] penicillin/streptomycin from the same companies as stated above. All cells were cultured in a 5% CO_2_ humidified incubator at 37 °C and controlled regularly for mycoplasma contamination. Throughout the experiments (unless otherwise stated), cells were treated with 1.5 µM AC220 (Selleckchem, S1526) and 10 µM spautin-1 (Sigma-Aldrich, SML0440) for CMA activation as previously described ^12^. The following compounds were used to treat cells in the indicated experiments: 50 µM CQ (Sigma-Aldrich, C6628), 50 nM Tranilast (Selleckchem, S1439), 10ng/ml TGFβ1 (PEPROTECH, 100-21).

### Cell proliferation and Clonogenicity

WT and KO cells of A549 and HT1080 cell lines were plated at a density of 3000 cells/well in their respective culture media in 6-well plates. After seeding, the number of cells (proliferation) was determined every day by using a hemocytometer and a light microscope.

To metabolically challenge the cells, they were grown in medium with either 10% galactose and 0.1 μM rotenone and cell numbers were counted every day for up to 5 days.

WT and LAMP2A KO cells were cultured in triplicate into 6 well-plates (200 cells per well) and allowed to form colonies over 8 days. Then, colonies were fixed with 1% paraformaldehyde and stained with 0.1% Crystal Violet solution. After washing and drying, the plates were imaged by a scanner (HP, G4010) and individual colonies were manually counted.

### Metabolomics

A targeted quantitative analysis was performed on LAMP2A WT and KO HT1080 xenografted tumors (25 mg tissue) using capillary electrophoresis mass spectrometry (CE-TOFMS and CE-QqQMS) in the cation and anion modes relative to internal standards at the Human Metabolome Technologies Inc. as described previously ^31-33^.

### Western blot

Cells were harvested and lysed using the RIPA lysis buffer supplemented with protease inhibitor cocktail (Roche, 04693124001). The frozen tumor tissues were homogenized with RIPA lysis buffer (with protease inhibitor) by a tissue lyzer (Retsch, MM400) and steel beads, and then centrifuged at 10 000 rpm for 10 min on a bench top centrifuge. The supernatant was transferred to a new tube. Protein concentrations in the cell or tumor lysates were determined using the bicinchoninic acid (BCA) assay (Thermo Fisher Scientific, 23225). After protein concentration adjustment the samples (with equal amounts of protein) were mixed with Laemmli loading buffer and boiled for 5 min at 95°C. Total protein extracts were separated in 12% acryl amide gels, transferred to nitrocellulose membrane and the membranes were blocked for 1 h in PBS with 5% skimmed milk and subsequently incubated overnight with the primary antibody at 4°C. After washing and incubation with the secondary antibody the recognized proteins were detected by enhanced chemiluminescence (ECL) substrate on X-ray film. The following antibodies were used in this study: LAMP2A (Abcam, ab18528, ab125068, ab240018), LAMP2B (Abcam, ab18529), LAMP1 (Santa Cruz, sc-20011), E-Cadherin (Cell Signaling, 24E10; Santa Cruz, sc71009), N-Cadherin (Cell Signaling, D4R1H; Santa Cruz, sc59987,), Vimentin (Cell Signaling, D21H3; Millipore, ab5733), TGFβR2 (Abcam, ab184948; Santa Cruz, E-6), TGFβR1 (Abcam, Ab235178), Smad2 (Cell Signaling, D43B4), Smad3(Cell Signaling, C67H9), Smad4 (Cell Signaling, D3M6U), pSmad2 (Cell Signaling, 138D4), pSmad3 (Cell Signaling, C25A9), HK2 (Thermo Fisher, 2H8L6), p53 (Santa Cruz, DO-1), ID-1 (BioCheck, BCH-1, clone 195-14), Vinculin(Abcam, ab129002,) and β-Actin (Santa Cruz, sc81178).

### Plasmids and siRNAs

The siRNAs targeting LAMP2A (65652 and 65654) were purchased from Shanghai GenePharma Co, Ltd. The siRNAs targeting TGFBR2 (s14077 and s14079) were purchased from Invitrogen. siRNA transfection was performed using the Lipofectamine 2000 Reagent (Invitrogen, 11668019) according to the manufacturer’s instructions. Plasmid pCMV6-XL5-LAMP2A was purchased from Origene (Cat nr #: SC118738) and used for transfection with ViaFect from Promega according to the manufacturers recommended procedure. The efficiency of siRNA and expression level of protein was monitored at 48– 72 h post-transfection by qPCR or Western blotting.

### cDNA synthesis and Real-time quantitative PCR (qPCR)

The total RNA was isolated from cells and tissues using the RNAqueous phenol-free total RNA isolation kit (Ambion, AM1912) according to the manufacturer’s instructions. A total of 1 µg RNA was used for cDNA synthesis with the IScript cDNA Synthesis Kit (Bio-Rad, 1708890), according to the manufacturer’s instructions. Approximately 200 ng of the cDNA sample was analyzed by quantitative PCR using Maxima qPCR SYBR Green Master Mix (Thermo Fisher Scientific, K0222) and amplified using the 7500 Real-Time PCR system (Applied Biosystems, Foster City, CA, USA). ACTB and HPRT were used as reference genes. ΔΔCt method was used to calculate the relative mRNAs levels after normalization with ACTB as a reference gene. The following primers were used:

LAMP2A (Fwd: 5’-ACTGTTTCAGTGTCTGGAGCAT-3’; Rev: 5’- ATGGGCACAAGGAAGTTGTC-3’);

LAMP2B (Fwd: 5’-AGGGTTCAGCCTTTCAATGT-3’; Rev: 5’- CTGAAAGACCAGCACCAACTA-3’);

LAMP2C (Fwd: 5’-TCAGTGTCTGGAGCATTTCAG-3’; Rev: 5’-GGTCAGAGTCAGCAGAACATT-3’);

LAMP1 (Fwd: 5’-TGTGGACAAGTACAACGTGAG-3’; Rev: 5’-CGTGTTGTCCTTCCTCTCATAG-3’);

E-Cadherin (Fwd: 5’-CTCGACACCCGATTCAAAGT-3’; Rev: 5’- CCAGGCGTAGACCAAGAAAT-3’);

Vimentin (Fwd:5’-GATTCACTCCCTCTGGTTGATAC-3’;Rev:5’-GTCATCGTGATGCTGAGAAGT-3’);

TGFβR2 (Fwd: 5’-ATGACATCTCGCTGTAATGC-3’; Rev: 5’-GGATGCCCTGGTGGTTGA-3’);

TGFβR1 (Fwd: 5’-GCAGAGCTGTGAAGCCTTGAGA-3’; Rev: 5’- TGCCTTCCTGTTGACTGAGTTG-3’);

β-Actin (Fwd: 5’-GCAAGCAGGAGTATGACGAG-3’; Rev: 5’- CAAATAAAGCCATGCCAATC-3’);

### CMA activity assay

HEK293FT cells were transfected with pSIN-PAmCherry-KFERQ-NE (Addgene, Cat# 102365), pLP1, pLP2 and pLP VSV-G in 10-cm cell culture dish for lentivirus production. Fresh medium (6 ml) was added to replace the used medium 24hrs after transfection. Two days later, medium was collected and filtered with 0.45 µm filter, then stored at -80°C for future viral transduction. To generate the stable FU-OV1 cells expressing PAmCherry-KFERQ-NE, FU-OV1 were seeded in 6-well plate. Next day, 1 ml of the lentivirus solution was added to the cells for transduction. 48h later, the medium was replaced by fresh medium containing puromycin (1μg/ml) for 5 days, whereafter cells were used for further analysis.

### Immunofluorescence

FUOV1 cells with stable expression of PAmCherry-KFERQ-NE were seeded on sterilized, round cover slips in 6-well plate. On the next day, the plate was placed in the UV chamber and exposed to UV light (365 nm) for 40 min to activate PAmCherry fluorescence. Then, cells were transferred back to the cell culture incubator and treated with either 50 nM Tranilast or 1.5 µM AC220 and 10 µM spautin-1. After treatment, cells were incubated with Hoechst 33342 solution for 10 min and washed three times with PBS. Afterwards, cells were fixed in 3% neutral buffered formalin on ice for 15 min, followed by three times washing in PBS and VECTASHIELD antifade mounting medium. The samples were measured under a Zeiss LSM700 confocal microscope. The puncta number was counted and analyzed from at least 60 cells under each condition.

### Assessment of mitochondrial oxygen consumption rates

Mitochondrial oxygen consumption rate (OCR) was measured in real-time using the XFp Extracellular Flux Analyzer (Seahorse Bioscience, MA, USA). Cells were seeded in 6-well plates at density 80 000 cells/well and treated with Tranilast (100 µM) for 6 hours. Then, the cells were transferred to an XFp miniplate at a density of 8000 cells/well in their respective growth media (supplied with same concentration of Tranilast for Tranilast-treated cells). On the next day, cells were washed twice with 100 µl of XF Base medium (Seahorse Bioscience), supplied with 1 mM pyruvate, 2 mM glutamine, and 10 mM glucose, and left in 180 µl of medium for 45 minutes incubation. Cells were subjected to analysis using XF Cell Mito Stress Test Kit (Seahorse Bioscience), following the manufacturer’s instructions. After baseline measurement, three following subsequent injections were made: 1 μM oligomycin (a mitochondrial ATP synthase inhibitor), 0.5 μM FCCP (a mitochondrial uncoupler), and 0.5 μM Rotenone/Antimycin A (Complex I/Complex III inhibitors). Wave (Agilent Technology) and GraphPad Prizm 9 software were used to analyze the obtained data.

### Analysis of G6PD enzymatic activity

G6PD enzymatic activity was determined using a kit (ab102529, Abcam, UK) according to manufacturer’s instructions. Cells were seeded on 10 cm Petri dishes and treated with 100 µM Tranilast for 24h. For each condition, 1×10^6^ cells were harvested, lysed in ice-cold PBS, and centrifuged at 12000 *x g* for 5 minutes at +4°C. The supernatant was collected for further analysis. The reaction mix for one reaction included: 46 µl G6PD Assay Buffer, 2 µl G6PD developer and 2 µl G6PD substrate. The measurements were performed at 450 nm wavelength on a GloMax Discover Microplate Reader (Promega, Madison, WI, USA) at 37°C within time range from 2 to 60 minutes. The enzymatic activities were normalized to protein concentrations, as determined by the BCA assay (Pierce, Thermo Fisher Scientific, Walthman, MA, USA).

### Statistics

Statistical analyses were performed using Prism v9.0. Data were presented as the mean ± S.D unless otherwise specified. Statistical tests are indicated in the corresponding figure legends. * P < 0.05, ** P < 0.01, *** P < 0.001.

## Acknowledgements

We thank Dr. Merve Kacal for generating the LAMP2A knockout of A549 and HT1080 cell lines and Professor Jorge Ruas, Dept. of Physiology and Pharmacology, Karolinska Institutet for comments on the manuscript. We are grateful to Dr. Alex Buko, Human Metabolome Technologies for the tumor metabolome analyses and Dr. Simon Stritt, Novogene for the RNA sequencing.

The work was supported by the following foundations: The Swedish Research Council (2021-01787) The Swedish Cancer Society (CAN 2017/466, CAN 2017/1015, 20 0979 PjF), The Wenner-Gren Foundations (UPD2020-0104 and UPD2021-0105), Stockholm Cancer Society (no: 174093), The Lars Hiertas Memorial Foundation and Stiftelsen Längmanska Kulturfonden.

## Authors contribution

H.V.N. and E.N conceived, designed, and supervised the study. XZ and YS performed most experiments with contributions from VS, EK, TB, BZ and VK. The animal experiments were designed by XZ, VS, HVN and MG and performed by XZ and VS. CL performed the immunohistochemical analysis of human tissues. SE and PH provided human lung cancer material.

EN and HVN wrote the manuscript with input from all authors.

## Figure Legends

**Figure S1.**
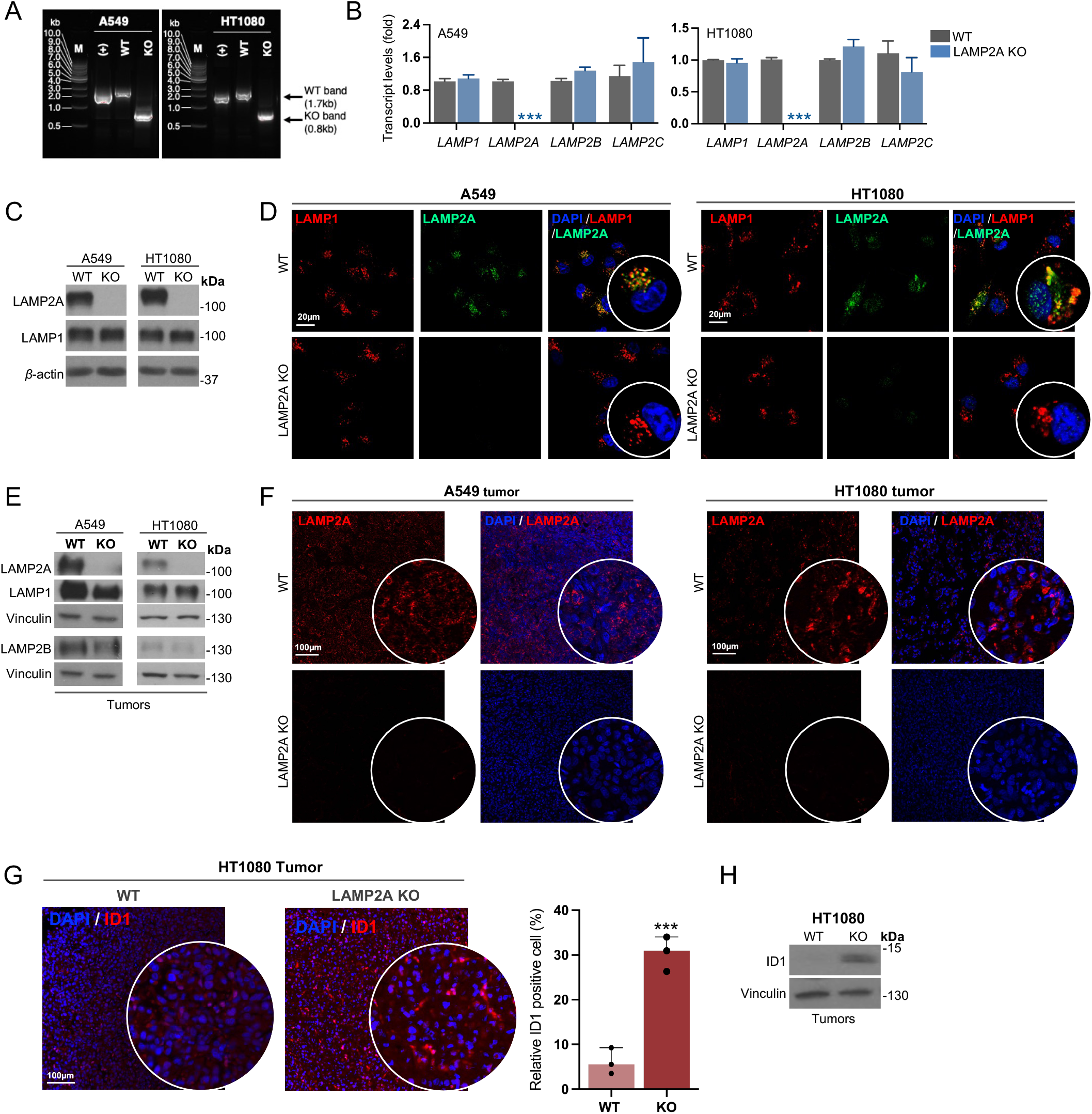
Generation and validation of an isoform-specific knockout of LAMP2A. **(A)** CRISPR/Cas9 genome editing and PCR genotyping of LAMP2A KO. **(B)** qRT-PCR analysis of *LAMP1, LAMP2A, LAMP2B* and *LAMP2C* expression in WT and KO cells from HT1080 and A549 cell lines **(C)** Western blot analysis of LAMP2A and LAMP1 in WT and KO **(D)** Confocal analysis of immunofluorescence with anti-LAMP2A (Green) and anti-LAMP1 (Red) antibodies in WT and KO cells from HT1080 and A549 cell lines. DAPI was used to counterstain cell nuclei. The scale bar (in white) is 20 µm. Insets at higher magnification as indicated. **(E)** Western blot analysis of LAMP2A, LAMP1 and LAMP2B in WT and KO xenograft tumors from HT1080 and A549 cell lines. Immunofluorescence staining for **(F)** LAMP2A or **(G)** ID1 in WT and KO HT1080 or A549 tumors. Scale bar is100 μm. Insets at higher magnification as indicated. Quantification of the ID1+ cell ratio per 20× field (bar graph). **(H)** Western blot analysis of ID1 in WT and KO xenograft tumors from HT1080 cell lines. Error bars, ±SD. ***p<0.001; two-tailed student’s *t*-test.

**Figure S2.**
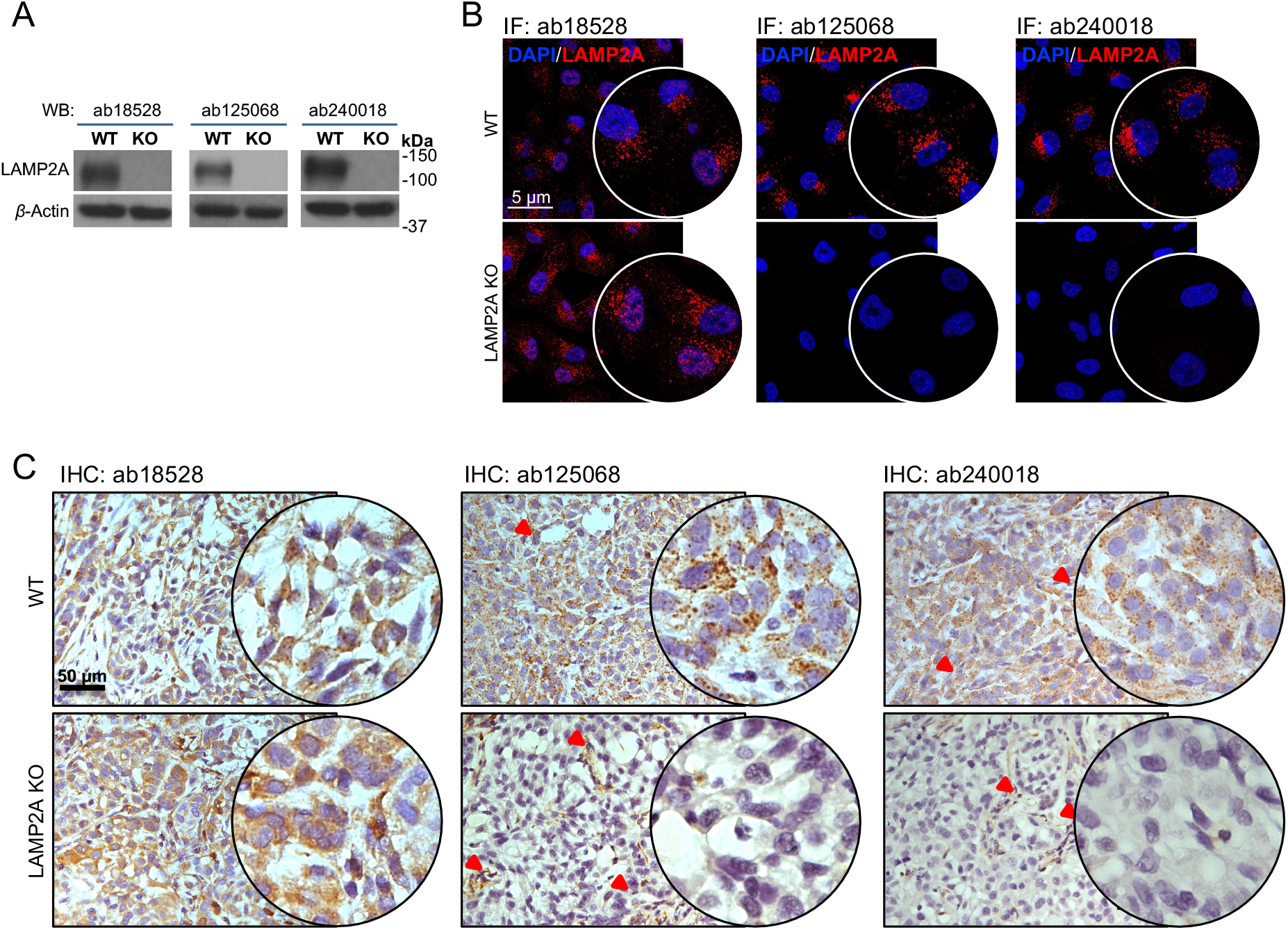
Antibody assessment of three commonly used LAMP2A antibodies. **(A)** Western blot analysis of three anti-LAMP2A antibodies in A549 WT and KO cells. **(B)** The immunostaining with three anti-LAMP2A antibodies in A549 WT and KO cells (red). DAPI was used to counterstain cell nuclei (Blue). The scale bar (in white) is 5 µm. Insets at higher magnification as indicated. **(C)** Immunohistochemical staining for LAMP2A in A549 WT and KO tumors by three LAMP2A antibodies (n = 3-4 mice). Red arrows mark the area of staining in the mouse stroma (tumor-supporting mice tissues). The scale bar (in black) is 50 µm. Insets at higher magnification as indicated.

**Figure S3.**
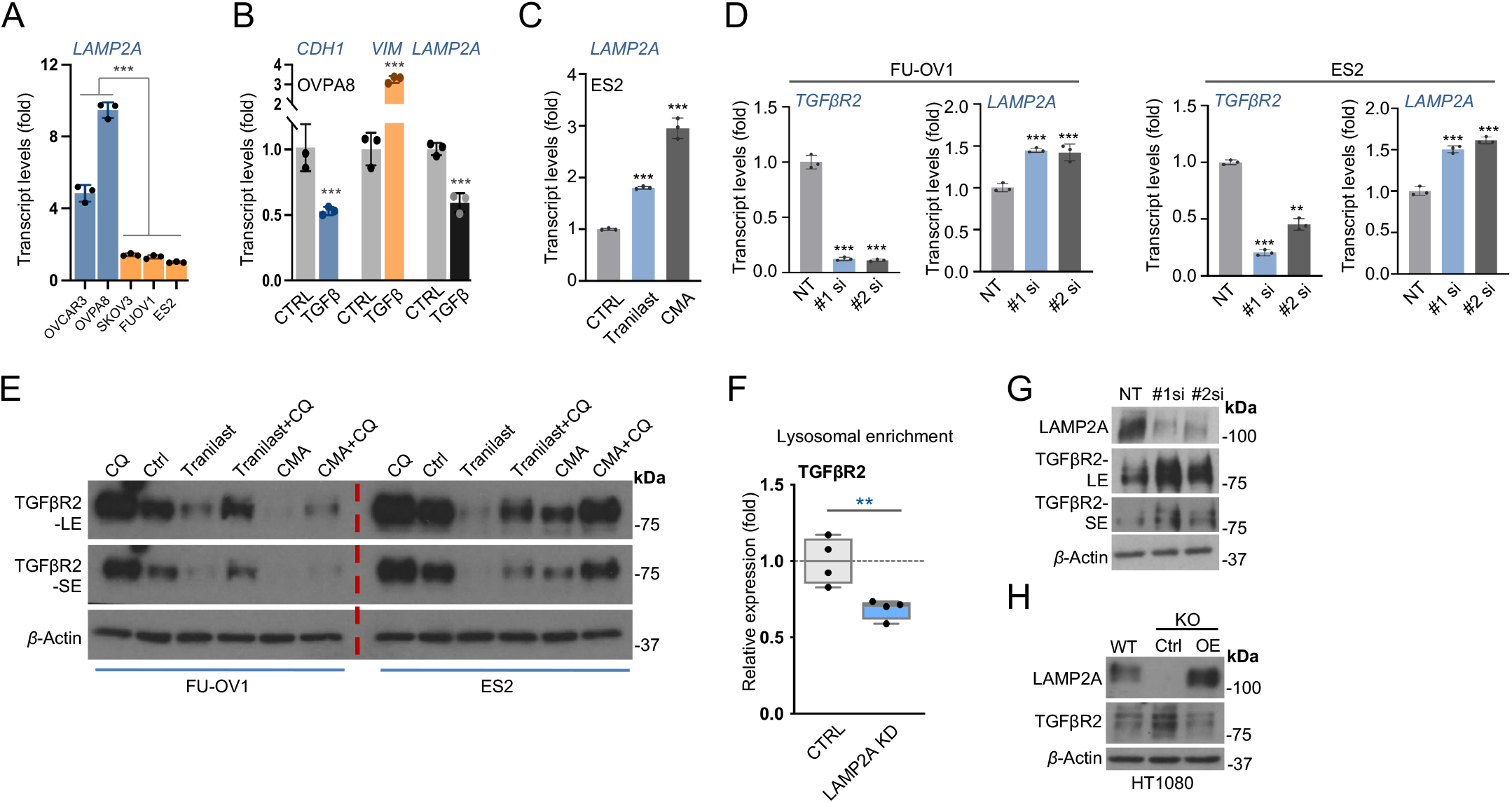
Mutual CMA and TGFβ signaling regulation. **(A)** qPCR analysis of LAMP2 expression in the indicated set of cell lines. **(B)** The effect of TGFβ treatment on *VIM, CDH1* and *LAMP2A* mRNA. The effect of TGFβ signaling inhibition (by Tranilast) or CMA activation on LAMP2A mRNA in OVPA8 cells. **(C)** The effect of CMA activation on TGFβR2 protein level in the ES2 cell line. **(D)** The effect of siRNA-mediated knockdown by two independent siRNAs targeting TGFβR2 on LAMP2A in FU-OV1 cells (left panel) and ES2 cells (right panel). **(E)** The effect of CMA activation, Tranilast, Tranilast +chloroquineon and CMA +chloroquineon treatment on TGFβR2. **(F)** The effect of LAMP2A knockdown on the TGFβR2 lysosomal enrichment as analyzed by mass spectrometry. **(G)** The effect of siRNA-mediated knockdown by two independent siRNAs targeting LAMP2A, on TGFβR2. **(H)** The effect of re-expression (RE) of LAMP2A in the LAMP2A KO HT1080 cells. Error bars, ±SD. **p<0.01, ***p<0.001; ns= non-significant; two-tailed student’s *t*-test.

**Figure S4.**
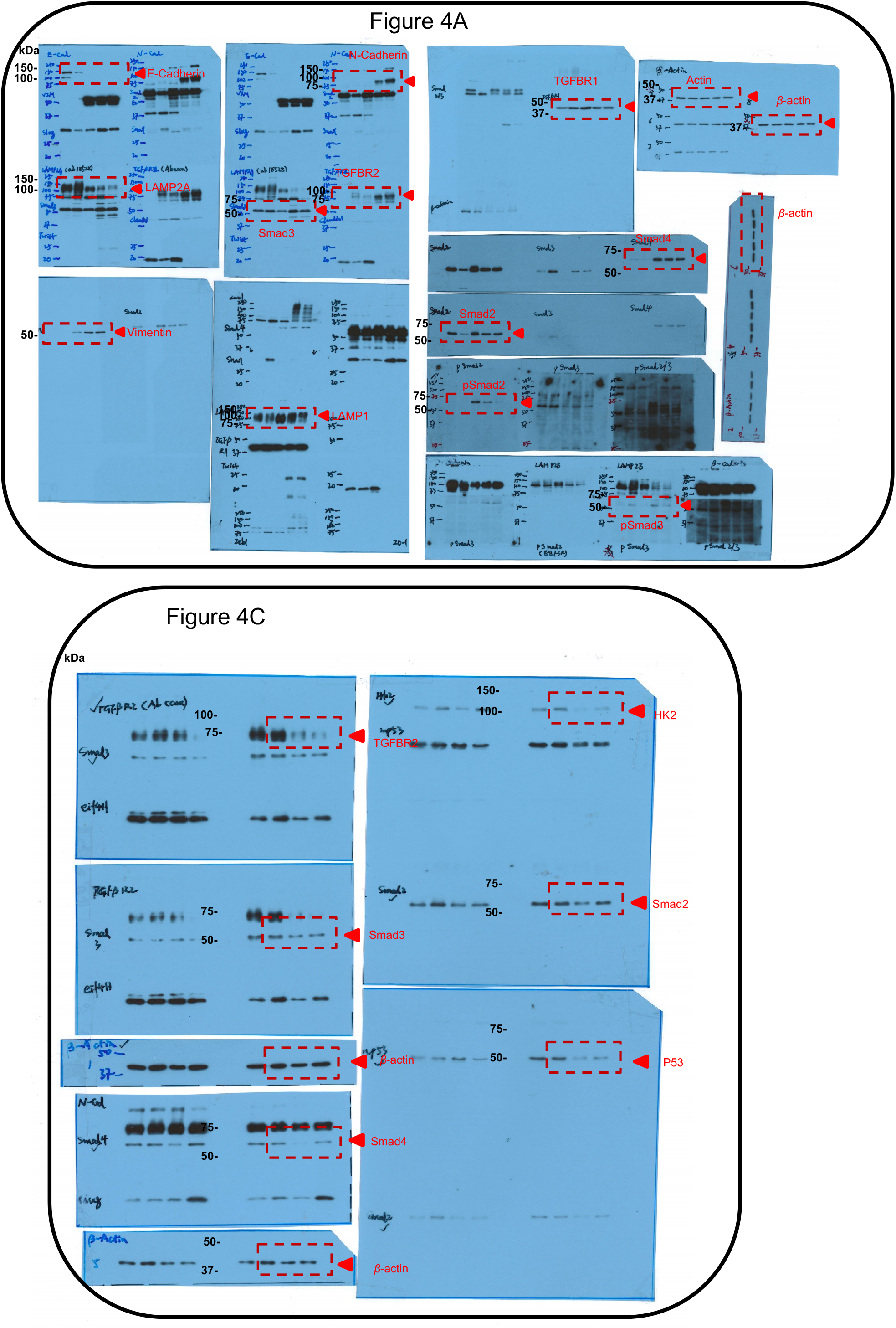
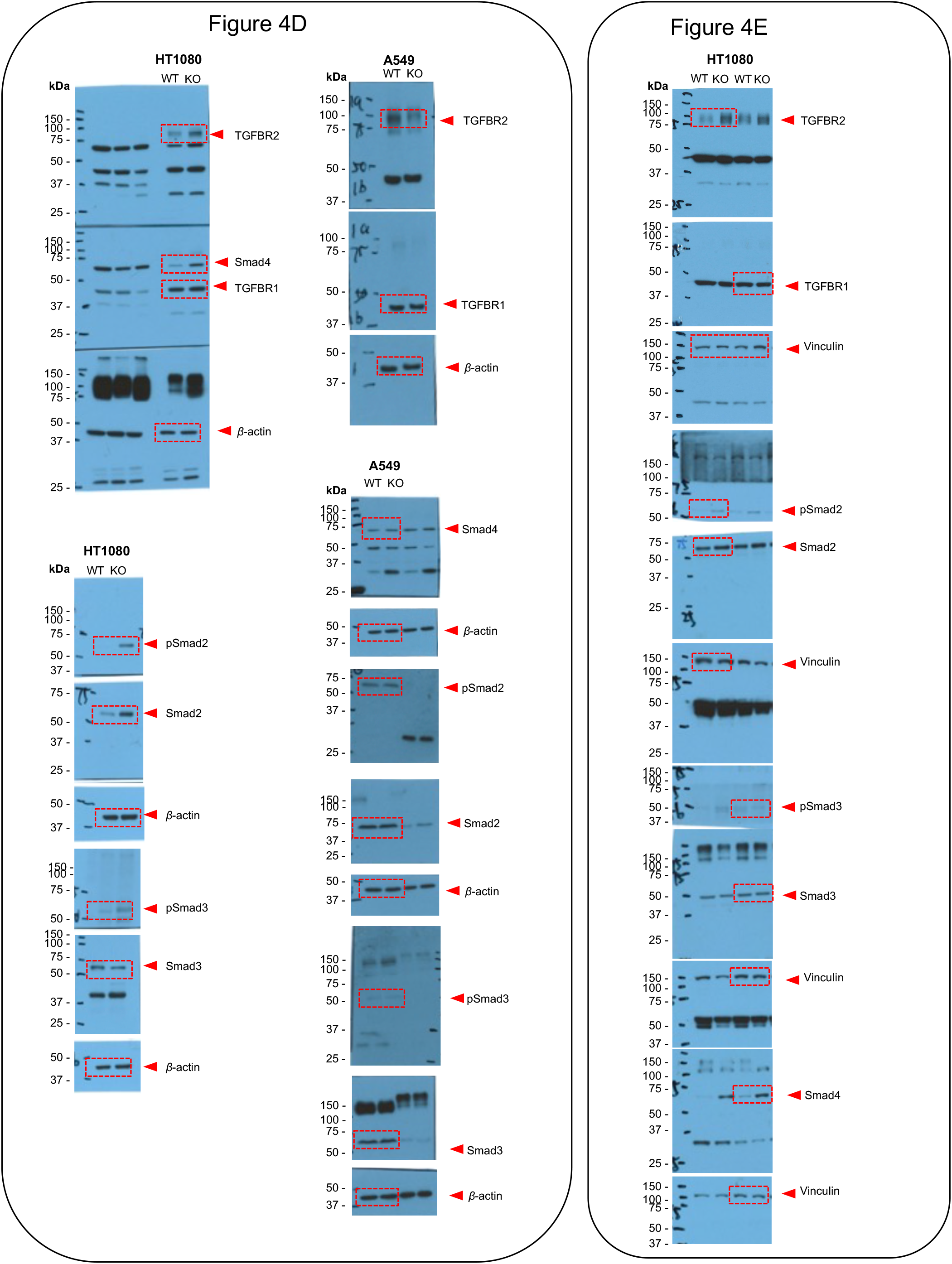
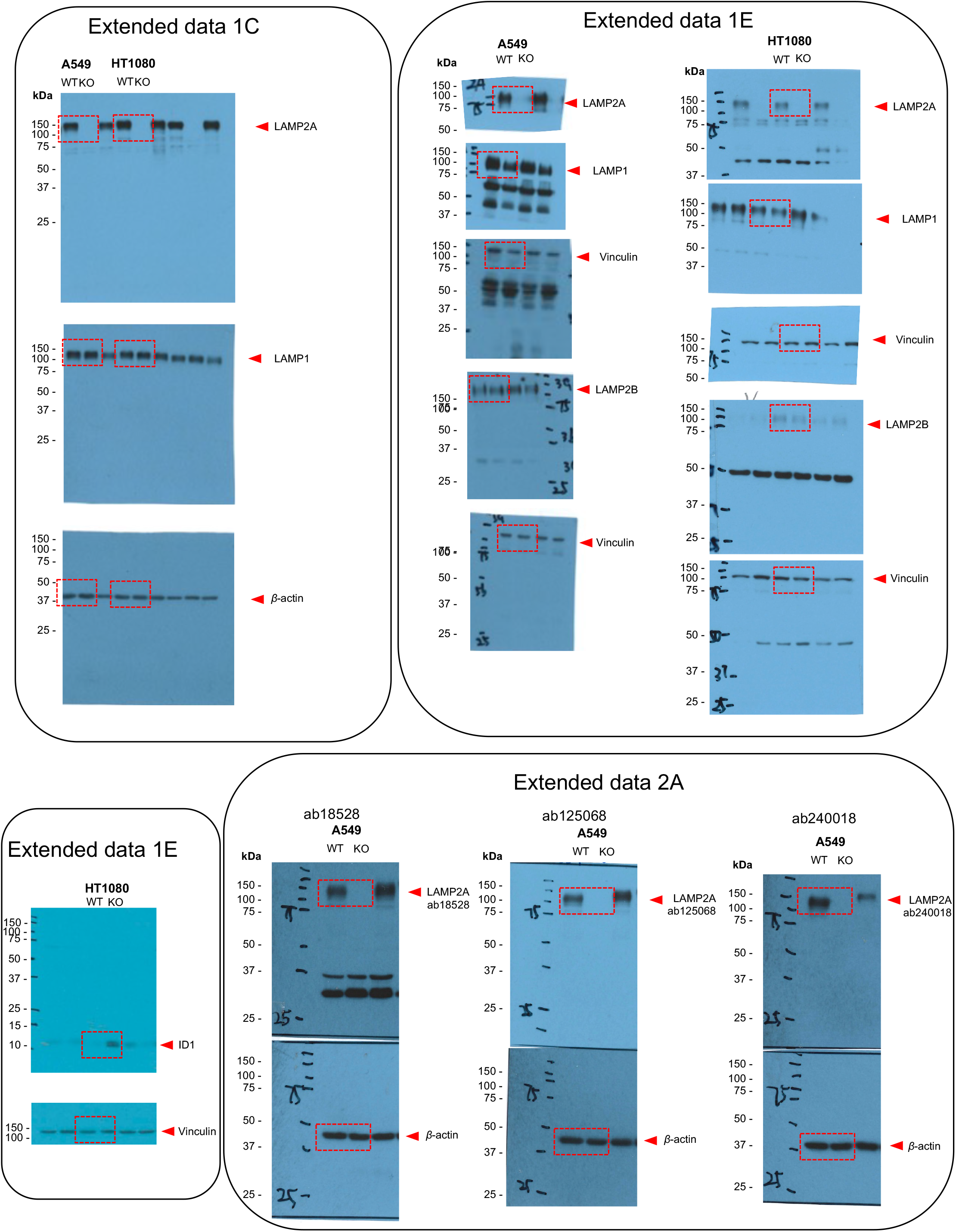
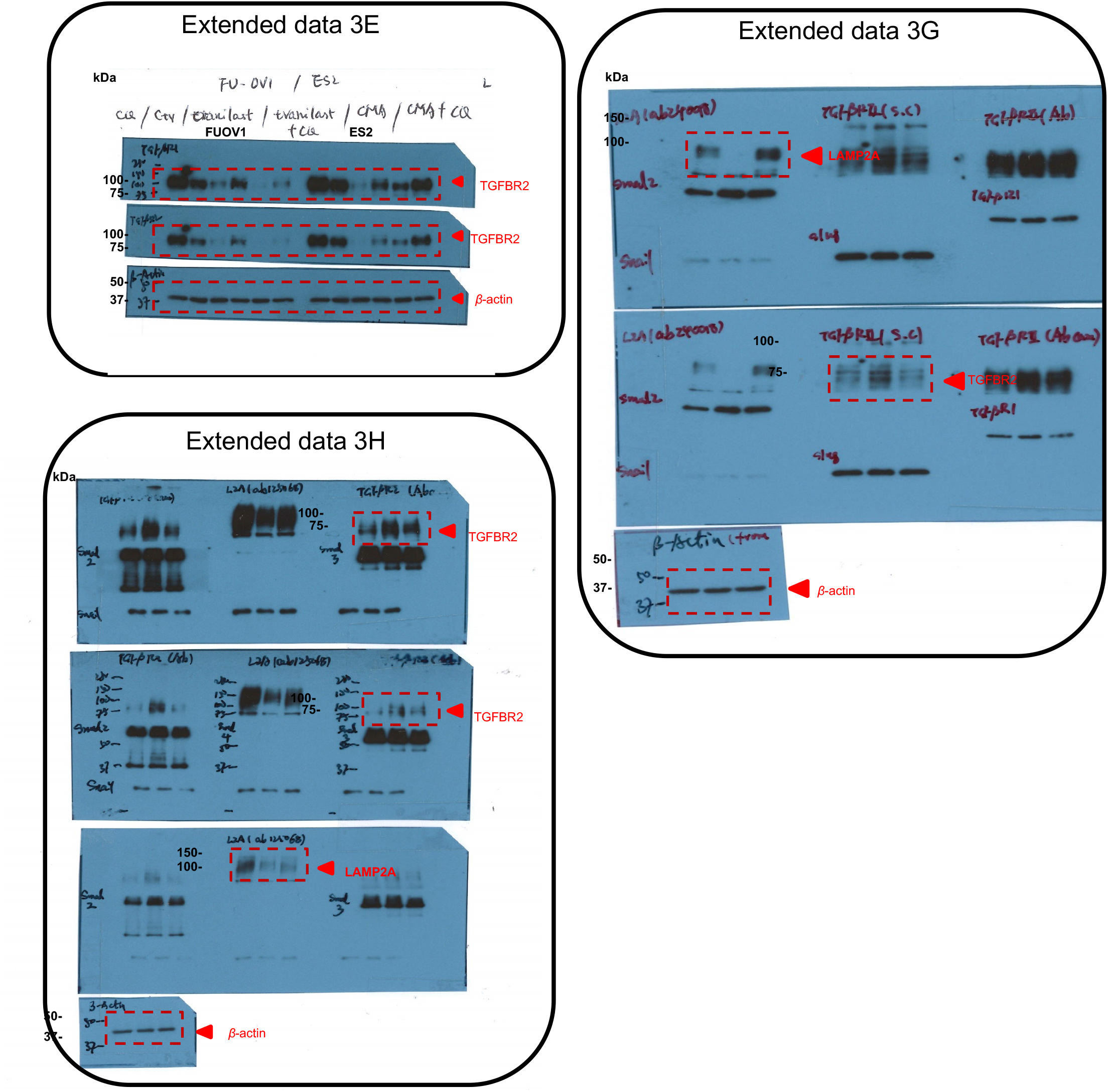
Original western blot scan for all figures.

